# Hippocampal Remapping as Hidden State Inference

**DOI:** 10.1101/743260

**Authors:** Honi Sanders, Matthew A. Wilson, Samuel J. Gershman

## Abstract

Cells in the hippocampus tuned to spatial location (place cells) typically change their tuning when an animal changes context, a phenomenon known as remapping. A fundamental challenge to understanding remapping is the fact that what counts as a “context change” has never been precisely defined. Furthermore, different remapping phenomena have been classified on the basis of how much the tuning changes after different types and degrees of context change, but the relationship between these variables is not clear. We address these ambiguities by formalizing remapping in terms of hidden state inference. According to this view, remapping does not directly reflect objective, observable properties of the environment, but rather subjective beliefs about the hidden state of the environment. We show how the hidden state framework can resolve a number of puzzles about the nature of remapping.

## Introduction

Place cells of the hippocampus fire when an animal occupies specific spatial locations (place fields; O’Keefe, 1976). Each place cell has its own respective place fields, so collectively the population of place cell comprise a map of an environment, in which each location corresponds to activity of a particular subset of place cells. The hippocampus is thought to use independent maps for each context. These independent maps can be observed through “place field remapping”, in which the location of a place field may change or the place field may disappear entirely between contexts (Muller and Kubie, 1987; Colgin et al., 2008). The sensitivity of place cells to context changes is consistent with many other studies implicating the hippocampus in context-dependent behavior (Holland and Bouton, 1999; Gershman et al., 2010; Smith and Mizumori, 2006b; Anagnostaras et al., 2001). Despite its acknowledged importance, the precise relationship between context changes and remapping has remained elusive, due in part to ambiguity as to what counts as context change.

Researchers have operationalized context in many different ways. For example, some researchers investigated the role of sensory cues (Knierim et al., 1998; O’Keefe and Conway, 1978; Muller and Kubie, 1987), whereas others investigated the effect of changing spatial location or geometry (Skaggs and McNaughton, 1998; Lever et al., 2002), or changing the task (O’Keefe and Speakman, 1987; Markus et al., 1995). Not surprisingly, different effects have been observed for these different manipulations, without cohering into a unified picture of how context changes determine remapping.

Some of the confusion about what counts as a context change is due to inconsistent definitions of the word “context”. Sometimes “context” refers to experimenter-defined variables, such as physical location or sensory cues. In other cases, “context” refers to the animal’s internal assessment of the environment as indicated by neural activity or behavioral response. For example, in the fear conditioning literature, animals are assumed to preferentially freeze in the “same” context as that in which they received the shock. This doesn’t necessarily have to be physically the same environment, as long as the animal infers that it is the same environment (Chang and Liang, 2017; Gershman et al., 2010). Invoking subjective inferential factors in the interpretation of remapping compels us to consider basic questions about the nature of these inferences. What is the animal’s hypothesis space? How does it represent and update beliefs over this hypothesis space?

The goal of this paper is to develop formal answers to these questions, and thereby provide a coherent account of diverse experimental findings. Key to this account is the idea that the relationship between observable properties of the environment (including context) and remapping is mediated by inferences about unobservable properties of the environment *(hidden states*). According to this view (see also Fuhs and Touretzky, 2007; Gershman et al., 2014; Penny et al., 2013), place fields remap when the animal believes that it has entered a new hidden state. By specifying the animal’s internal model of how hidden states relate to observable stimuli, we can make principled predictions about when, why and how place fields remap.

Before describing the details and applications of this computational framework, we will briefly review some of the key empirical and theoretical background.

## Empirical background

Remapping phenomena have been divided into several classes (Colgin et al., 2008; Muller, 1996). At the extremes, there is “global” or “complete” remapping (where no place fields are shared between contexts) and “null” or “lack of” remapping (where all place fields are shared between contexts). Between these extremes is “partial remapping” (where some place fields are shared between contexts but some are not) and “rate remapping” (where place fields are shared between contexts but have characteristically different firing rates). However, none of these categories can be regarded as strictly exclusive.

The extent to which place fields are shared between contexts can be quantified by looking at the spatial correlations of place cell firing rates between contexts. Although studies report correlations near zero between place fields in different contexts (Leutgeb et al., 2004; Muller and Kubie, 1987; Schlesiger et al., 2015), there are reasons to believe that correlations are not actually zero. A recent report suggests that previous observations of global remapping might be artifacts of misalignment of maps between contexts (Kinsky et al., 2018). Some place cells have been found to consistently encode reward across virtual reality contexts that otherwise express “global remapping” (Gauthier and Tank, 2018), so there is at least one class of place cells that have recently been found not to remap across contexts. More generally, many studies reporting global remapping report low but non-zero correlations (Leutgeb et al., 2004, 2005b; Skaggs and McNaughton, 1998; Spiers et al., 2015).

Conversely, studies reporting lack of remapping never report perfect place field overlap between contexts. Indeed, even within a single context, patterns of spatial firing show variability over time, as if more than a single map is used in a given context (Fenton and Muller, 1998; Kay et al., 2019; Kelemen and Fenton, 2016). Additionally, the extent of remapping for repeated presentations of the same context depends on the amount of experience the animal has had (Law et al., 2016).

Rate remapping is also not a strict category. Manipulations used to generate rate remapping do so for a fraction of the place cell population, while other cells in the population maintain or lose their place fields (Wood et al., 2000; Leutgeb et al., 2005b). In this way, rate remapping is always accompanied by partial remapping. Additionally, protocols for generating rate remapping can sometimes produce a range of remapping states during learning, ranging from no remapping to global remapping. For example, Leutgeb et al. (2005b) found rate remapping when comparing place field maps between circle and square enclosures. However, Lever et al. (2002) make the same comparison between circle and square enclosures, and find rate remapping as an intermediate state as the animal transitions from no remapping to global remapping over the course of learning.

The complications discussed above highlight the fact that virtually all remapping is partial remapping. Place cell responses to manipulations are extremely heterogeneous (Lee et al., 2004; Shapiro et al., 1997; Chen et al., 2013; Anderson and Jeffery, 2003). Additionally, remapping behavior can vary across animals (Wills et al., 2005; Lever et al., 2002) as well as across laboratories (Guzowski et al., 2004; Wills et al., 2005; Leutgeb et al., 2005a; Colgin et al., 2010, see the “Morph Experiments” section of the Results for an in-depth exploration of one example). We will argue that this heterogeneity arises from variability in beliefs across animals.

## Theoretical background

Our theory of hidden state inference is motivated by, and builds upon, prior research into the nature of context-dependent learning. Since Pavlov, experimentalists have recognized that extinguishing an association after Pavlovian conditioning is not the same as unlearning it. The association can return under a variety of circumstances (Bouton, 2004), such as returning the animal to the conditioning context, or simply waiting a period of time before testing the animal. These phenomena seem to suggest that the animal is forming a new memory during extinction, which could compete with the conditioning memory at the time of retrieval. Context, on this view, serves as a particularly powerful retrieval cue. The fundamental challenge posed by this interpretation is to define precisely the conditions under which a new memory is formed or an old memory is updated, and the conditions under which a particular memory is retrieved at the time of test.

One approach to these questions is to frame them in terms of hidden state inference (Gershman et al., 2010, 2017): new memories are formed when an animal has inferred that it has encountered an unfamiliar (previously unvisited) state, and old memories are updated when it has inferred that it has encountered a familiar state. As we formalize below, these inferences can be calculated using Bayes’ rule, which computes a posterior probability distribution over hidden states by integrating prior beliefs about the hidden states with the likelihood of those hidden states given the animal’s observations. The hidden states are sometimes interpreted as *latent causes* (Courville et al., 2006; Gershman and Niv, 2012), to emphasize the idea that the animal is forming beliefs about the causal structure of the environment.

The state inference framework can naturally explain many animal learning phenomena (see Gershman et al., 2015, for a review). For example, a conditioned response takes longer to extinguish when reward is delivered probabilistically during the acquisition phase, a phenomenon known as the *partial reinforcement extinction effect* (e.g., Gibbon et al., 1980). This phenomenon is surprising for classical associative learning accounts, since the learned association should be weaker under partial reinforcement, and hence should be *faster* to extinguish. According to the state inference framework, partial reinforcement renders the hidden state ambiguous; it takes more extinction trials until the animal is confident that acquisition and extinction trials were generated by different states (Courville et al., 2006; Gershman and Blei, 2012).

In this paper, we argue that the same framework can unify many different place field remapping phenomena, under the assumptions that (i) each map corresponds to a unique hidden state, and (ii) a map is activated in proportion to the posterior probability of the corresponding hidden state. A closely related idea was pursued by Fuhs and Touretzky (2007), to which we owe the inspiration for the present work. Our goal is to explain a significantly broader range of phenomena using a somewhat simpler model, and to resolve a number of lingering empirical puzzles.

## Results

### Conceptual overview of the model

The computational problem facing the animal is to infer the posterior probability of each hidden state *c* given its observations **y** (e.g., geometric or color features of a box), as stipulated by Bayes’ rule:

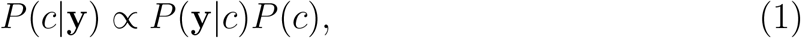

where *P*(**y**|*c*) is the likelihood of the observations under the hypothetical state *c*, and *P*(*c*) is the prior probability of state *c*. A more detailed formal description of these terms can be found in the Materials and Methods. In this section, we describe intuitively what they mean and how they work.

The animal is presented with observations that are generated by an unknown number of states through a process that the animal is not aware of (left side of Fig. 1A). The animal builds an *internal model* of the world (thought bubble in Fig. 1A). That model doesn’t have to mimic the world exactly, it simply needs to be flexible enough to be able to capture the structure that it is presented with. We suggest that the animal’s internal model provides a generative “recipe” through which it assumes observations are produced: first a state is sampled from *P*(*c*), and then an observation is sampled from the distribution associated with that state *P*(**y**|*c*). The job of the animal is to invert this generative process and infer the posterior probability of each hidden state *c* given its observations **y**. Since different states could theoretically produce the same observations, the animal is faced with fundamental ambiguity. The posterior distribution *P*(*c*|**y**) represents the animal’s uncertainty about the hidden state. As it collects more observations and thereby reduces its uncertainty, the posterior will tend to progressively concentrate on a single explanation of which observations come from which states.

Because there is no reason to assume that the animal has *a priori* knowledge about the set of states, we allow the state space to potentially grow as the animal collects new observations. The animal starts off with a single state, and at each new observation it can assign some probability to a new state or one of its previously inferred states. As detailed in the Materials and Methods, we accomplish this using a Bayesian nonparametric prior over hidden states. Importantly, this prior favors a small number of hidden states, encoding a form of “simplicity bias” or Occam’s razor.

As mentioned in the Introduction, we assume a one-to-one correspondence between hidden states and maps. Thus, we transpose the question “did the place field remap?” to “were these observations generated by the same hidden state?” More precisely, we report the log posterior probability ratio between 1-state and 2-state hypotheses (or *evidence ratio*, for brevity), which we take to be related to the degree of remapping (see Materials and Methods for definitions of two versions of the evidence ratio: the partition evidence ratio and the state evidence ratio). When the evidence ratio is near 0, the animal is indifferent between the two hypotheses, and in this case we expect partial remapping. No remapping occurs when the evidence ratio is strongly positive (favoring the 1-state hypothesis), rate remapping occurs when the log probability ratio is weakly positive, and global remapping occurs when it is strongly negative. Keep in mind, following our overview of the literature in the Introduction, that these are heuristic categories without strict boundaries. On the probabilistic view, these categories occupy different points along a spectrum.

To develop some intuitions for how the model works, we begin with a toy example. We generate observations from a 2D Gaussian with mean *μ* = [0,0] and a diagonal covariance matrix with standard deviations *σ* = [1,0.2], such that the distribution is more spread out along feature 1 than along feature 2. Figure 1B shows a single observation drawn from that distribution (black x). Conditional on this observation, we can measure, for each point in the feature space, the state evidence ratio (i.e., the relative probability that a particular point would be assigned to the same hidden state as the previous observations as compared to a novel hidden state). The state evidence ratio is shown as a heat map in Fig. 1B. Points close to the original observation are more likely to be assigned to the same state as the original observation. Overall, the model is relatively uncertain, reflected in the fact that the state evidence ratio is near 0 for all points in the panel.

**Figure 1:**
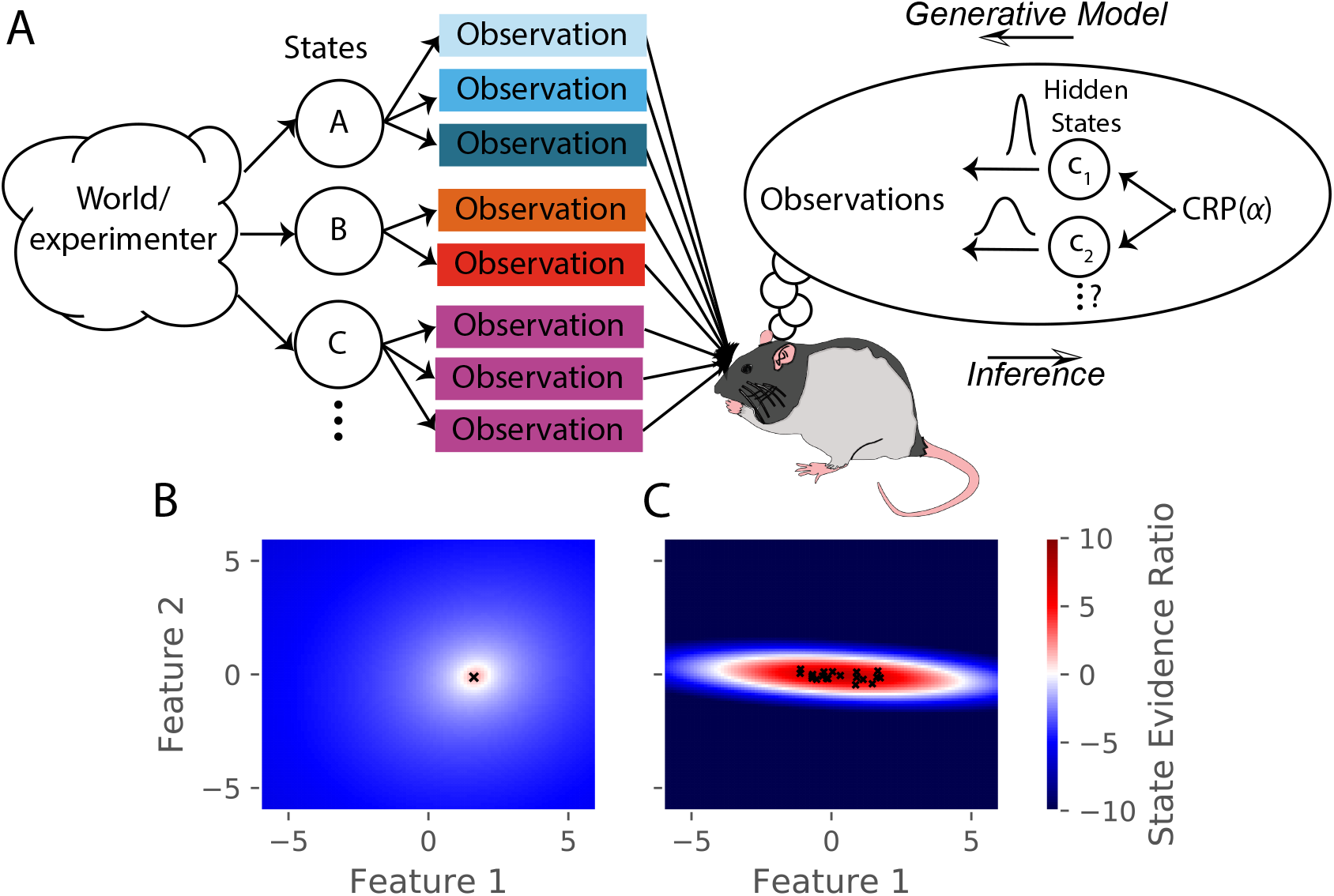
(A) Schematic of hidden state inference. We impute an internal generative model to the animal, according to which observations are generated by a small number of hidden states. States are sampled from the Chinese Restaurant Process, parametrized by *α* (see Materials and Methods for details). Each state is associated with a particular distribution over observations. The animal receives those observations but does not have direct access to the states that generated them. We model the animal as probabilistically inverting this generative model by computing the posterior distribution over hidden states given observations. (B-C) Demonstration of inference process. Observations are black x’s, background color is the relative probability of assigning a new observation to same hidden state as black x’s (as opposed to assigning it to a novel hidden state) after (B) 1 observation and (C) 20 observations.

The probability of assigning novel observations to the same state as past observations changes over the course of learning. Computing the state evidence ratio after the model has seen 20 draws from the same distribution (black x’s in Fig. 1C), we see that the model has increased certainty: the probability of assigning a point to the same hidden state has a higher central peak and falls off more quickly with distance from the previous observations. Moreover, the model has learned the shape of the generative distribution.

This example showcases a trade-off of hidden state inference: it can capture the shape of the data, but requires experience in order to do so.

### The effect of sensory cues

One of the first questions asked about hippocampal remapping was which sensory cue controls whether a map is used. The first study of remapping (O’Keefe and Conway, 1978) found that in an environment with four cues, some place fields disappeared with the removal of one or two cues, but most place fields maintained their firing with the removal of any two cues. In more modern terms, removal of a subset of cues caused partial remapping, but there was not a one-to-one correspondence between place fields and cues. Thus, from the very beginning it was clear that remapping is not in response to cues but in response to cue constellations (see also Shapiro et al., 1997; Fenton et al., 2000; Muller and Kubie, 1987). Each of these studies involved separately rotating or removing groups of stimuli, finding that many place fields that rotated when a given stimuli was rotated still maintained their firing when that stimuli was removed. A similar early result was that of O’Keefe and Speakman (1987), where cues necessary for orientation of the map were removed, but the place cell map was maintained. The significance of these results is that the place field map is responsive to cues but is not controlled by cues in a one-to-one fashion.

Viewing remapping as hidden state inference provides an important insight into this behavior. Our model posits that the cues jointly inform the posterior over hidden states. Individual cues will typically only exert a weak effect on the posterior, and hence exert only a weak effect on remapping.

To simulate the effect of cue configurations on remapping, we assume that the observation vector consists of four features, each drawn from a Gaussian with mean 0 and standard deviation of 0.3. We provide the model with 20 observations drawn from that distribution and then provide one of four probe observations. For each probe, we compute the state evidence ratio (Fig 2).

The first probe is an observation where each feature has a value of 0 (no cues changed). The model prefers assigning the probe observation to the same hidden state as the previous observations, corresponding to no remapping. The second probe is an observation where the first feature has a value of 1 and the other features have a value of 0 (cue 1 changed). The third probe is an observation where the first and last features have a value of 1 and the other features have a value of 0 (cues 1 and 4 changed). For both of these, the model produces an evidence ratio near 0, registering a high level of uncertainty about the hidden state (i.e., partial remapping). The fourth probe is an observation where all four features have a value of 1 (all cues changed), for which the model prefers assigning the probe observation to a new hidden state, corresponding to global remapping. These simulations demonstrate how the model is sensitive to the configuration of cues; no one cue completely controls remapping, consistent with the experimental data reviewed above.

Another aspect of these simulations worth highlighting is the fact that they are probabilistic. The representation of uncertainty in hidden state identity corresponds in an important way with the result that hippocampal maps during two experiences are almost never entirely overlapping nor entirely independent. From the perspective of our model, this “partial remapping” reflects the inherent uncertainty about whether different observations are drawn from the same distribution.

### Experience-dependent remapping

The previous section addressed the study of how sensory cues control place field remapping. Another line of research has studied how more diffuse contextual cues control remapping, but the answer was invariably that it depended on prior experience (Knierim et al., 1995; Sharp et al., 1990; O’Keefe and Speakman, 1987; Breese et al., 1989; Knierim et al., 1998; Bostock et al., 1991; Shapiro et al., 1997). One prime example of this is the role of environmental geometry (the shape of the recording arena). Initially, it was thought that different geometries necessarily corresponded to different maps (Muller and Kubie, 1987; Quirk et al., 1992), but recordings had always been done in familiar environments. The first group to record throughout the course of learning found that there was no consistent relationship between environment shape and inferred hidden state (Lever et al., 2002). In this experiment, place cells were recorded in rats who were alternately placed in square and circle boxes occupying the same location in the recording room day after day. Early in learning, there was limited remapping. Only after extensive experience in the two boxes did the animals remap between the two boxes (Fig. 3A). This indicates that the sensitivity to context changes changes with experience. Analogous results have been found for the effects of experience on remapping in response to other manipulations (Bostock et al., 1991; Shapiro et al., 1997). These effects are hard to explain in terms of fixed contextual boundaries governing remapping. It is naturally explained by the hidden state inference perspective, which posits that uncertainty about hidden states evolves as more data are observed. In particular, distinctions between hidden states are acquired gradually, such that substantial remapping should only be observed after extensive experience, counteracting the “simplicity bias” favoring a small number of hidden states.

**Figure 2:**
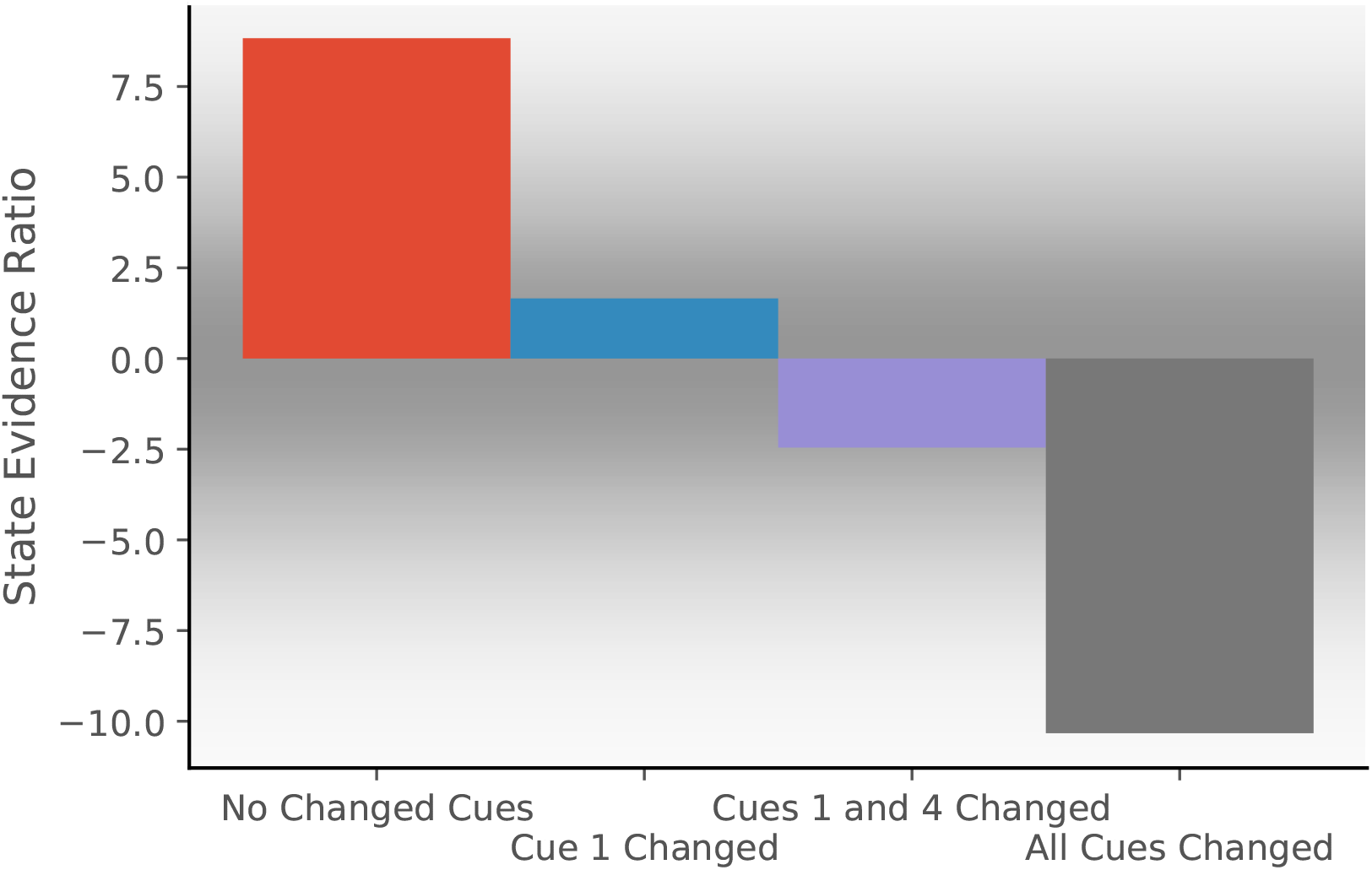
Observations are generated from a distribution with four features, each drawn from a Gaussian with mean 0 and standard deviation of 0.3. We train the model with 20 observations drawn from that distribution. We then compare the posterior probability of assigning a probe observation to the same hidden state as the previous observations vs. assigning it to a novel hidden state (Eq. 10 for same *c* vs. novel *c*). The first probe is an observation where each feature has a value of 0 (no cues changed). The model prefers assigning this probe observation to the same hidden state as the previous observations, corresponding to no remapping. The second probe is an observation where the first feature has a value of 1 and the other features have values of 0 (cue 1 changed). The third probe is an observation where the first and last features have a value of 1 and the other features have values of 0 (cues 1 and 4 changed). For both of these, the model assigns a state evidence ratio near 0, representing relatively high uncertainty about hidden state assignment, which corresponds to partial remapping. The grey background has saturation proportional to a Gaussian centered at 0 with a standard deviation of 5; values with a grey background can be heuristically thought of as partial remapping, whereas values with a white background can be thought of as either complete remapping or lack of remapping depending on whether 2 states are more likely (negative values) or 1 state is more likely (positive values). The fourth probe is an observation where all four features have values of 1 (all cues changed), for which the model prefers assigning the probe observation to a new hidden state, corresponding to global remapping.

We simulate these experiments qualitatively in the following way. We take observations to be 1D for simplicity, where the single dimension is the feature along which the distinction is learned. For example, in the circle-square experiment (Lever et al., 2002), the dimension would be the shape of the enclosure. We generate observations from two Gaussians (corresponding to the circle and square contexts) with *μ*_1_ = –0.5, *μ*_2_ = 0.5, *σ*_1_ = *σ*_2_ = 0.1 (Fig. 3B). We alternate drawing observations from each distribution. After each pair of draws, we compute the partition evidence ratio (in this case, the relative probability of the hypothesis that all observations up to that point were drawn from a single hidden state against the hypothesis that all observations up to that point had been drawn from two alternating hidden states).

Early in training, there is uncertainty about how many hidden states there are (Fig. 3C); the evidence provided by the observations is not yet sufficiently strong to overwhelm the simplicity bias of the prior. As more data are observed, The two-state hypothesis is eventually favored over the one-state hypothesis. The hidden state inference perspective thus explains why context-dependent remapping only emerges gradually with experience.

### Stabilization of maps over time

Maps take time to stabilize: repetition of a novel environment induces less map similarity than repetitions of a familiar environment (Frank et al., 2004; Leutgeb et al., 2004; Law et al., 2016). In particular, Law et al. (2016) alternated presentation of two environments. They found that intra-environment map similarity went up as a function of experience (Fig. 4A). These results are difficult to explain under the assumption that remapping is induced by the discrepancy between expectations and current cues exceeding a fixed threshold (Jeffery, 2003). Long-term potentiation (LTP) had been tied to map stabilization (Kentros et al., 1998; Cobar et al., 2017), but the speed with which LTP can create place fields (single trials; Bittner et al., 2017) is inconsistent with the slowness of map stabilization. The hidden state inference perspective offers a different interpretation of map stabilization: as an animal gains more experience with a particular state, it sharpens its representation of that state (i.e., its uncertainty about the distributional statistics decreases), and consequently it becomes more confident in recognizing repetitions of that state.

**Figure 3:**
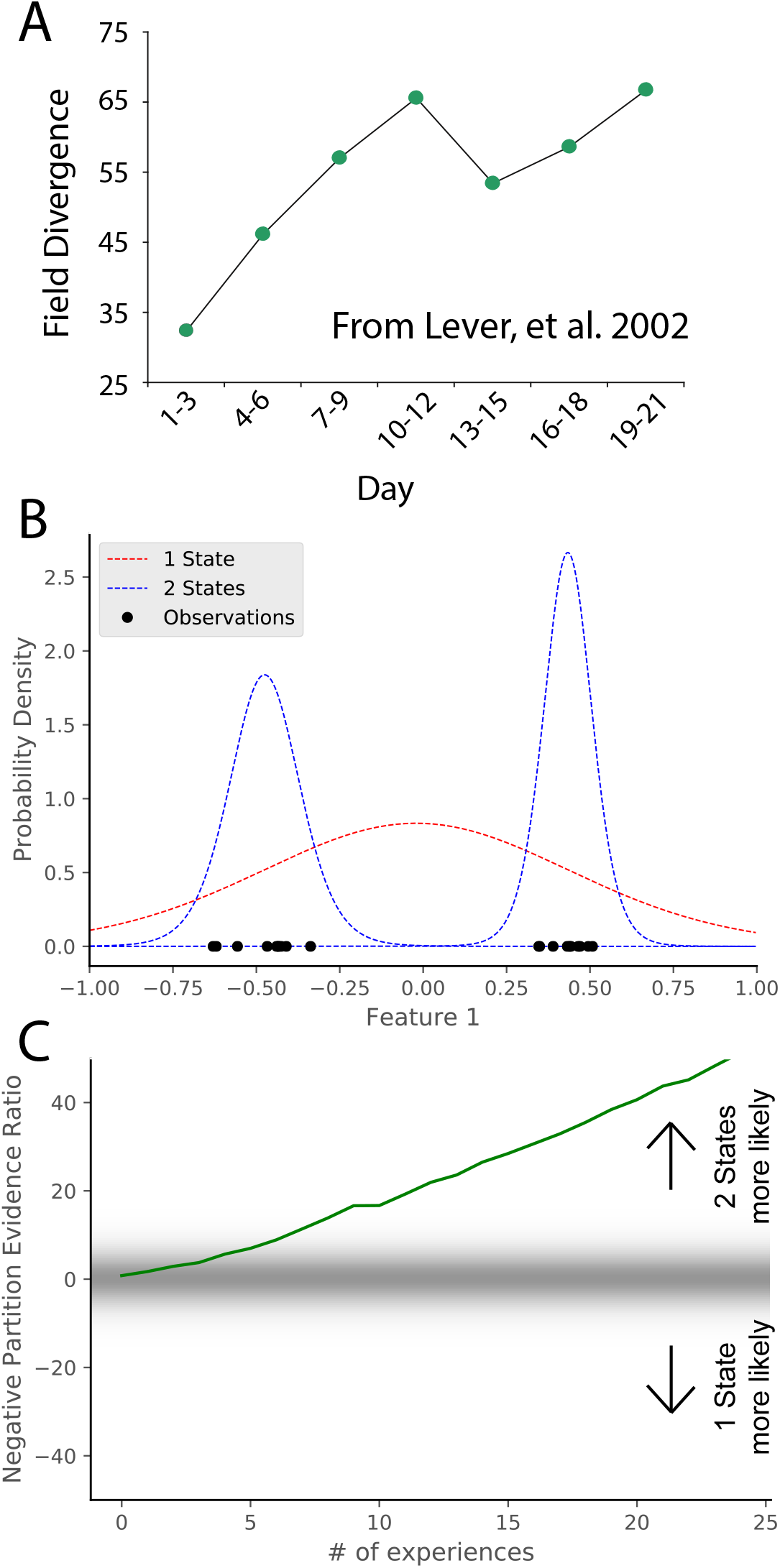
(A) Data from Lever, et al. (2002), who compared place cell representations between alternating presentations of square and circle boxes. Field Divergence is expressed in percent and represents the fraction of place fields that remap between the two enclosures. The representations of the enclosures are initially similar, but diverge with learning. (B) Simulated observations (black dots) are generated from Gaussians centered at −0.5, 0.5. The model compares the posterior probability of the observations coming from 1 inferred hidden state (red) or 2 inferred hidden states (blue). (C) The relative probability assigned to the observations coming from 2 hidden states vs. 1 hidden state (Eq. 7) is shown as a function of amount of experience. Early on, there is uncertainty about how many hidden states there are, whereas later 2 hidden states is more probable, similar to the empirical observations. As in Fig. 2, values with a grey background can be thought of as partial remapping whereas values with a white background can be thought of as either complete remapping or lack of remapping depending on whether 2 states are more likely or 1 state is more likely. Note that the axis here has been flipped relative to Fig. 2 in order to match the axis of the empirical results shown in panel A.

**Figure 4:**
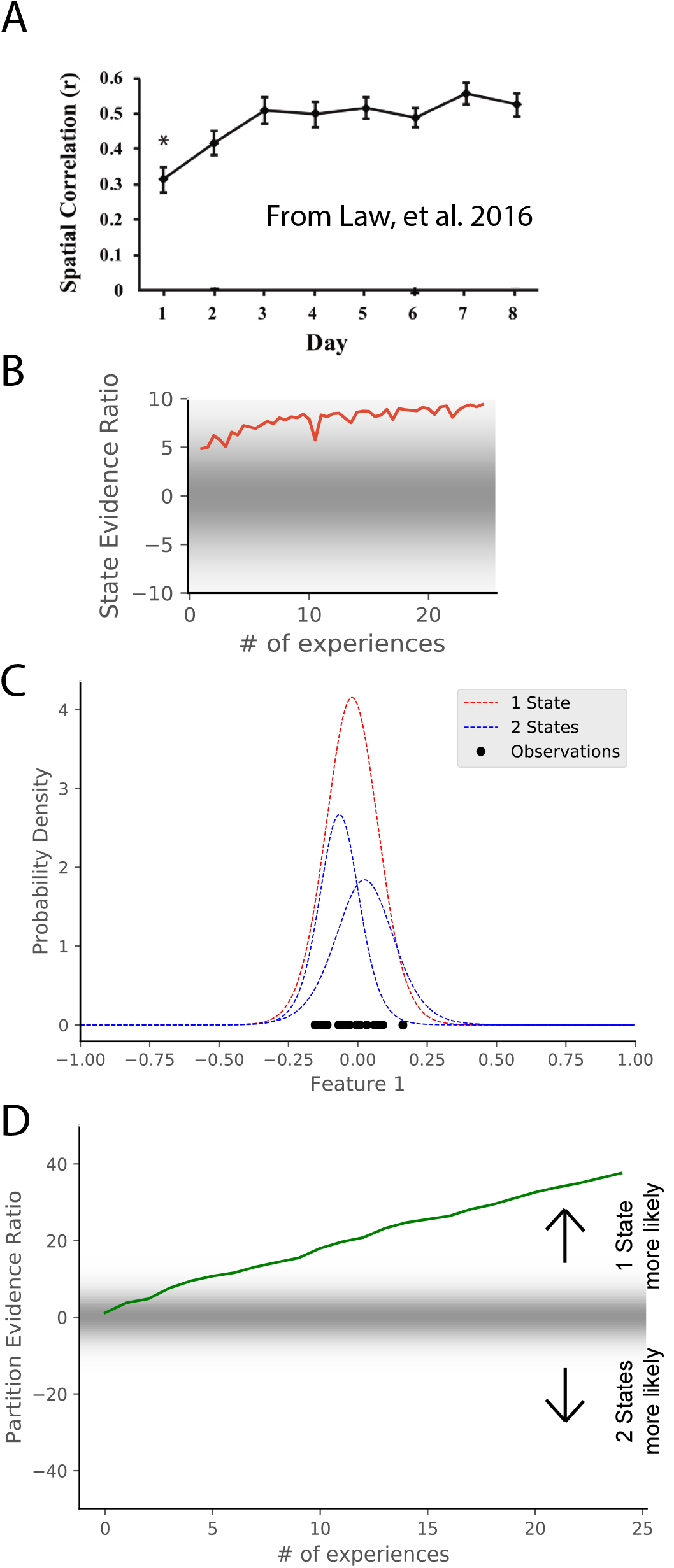
(A) Data from Law et al. (2016), showing the spatial correlation of the hippocampal map in repeated presentations of the same environment over multiple training days. Initially, the correlation is low, indicating extensive remapping between observations, but over the course of training the extent of remapping between observations decreases. (B) For each observation from the simulations in Fig. 3, we calculate the relative probability of assigning that observation to previous observations from that hidden state compared to coming from a novel hidden state (Eq. 10). The probability associated with the novel hidden state (and transitively, with remapping) decreases over the course of training. (C) Observations (black dots) are generated from two Gaussians, both of which are centered at 0. The model compares the posterior probability of the observations coming from 1 inferred hidden state (red) or 2 inferred hidden states (blue). D) The relative probability assigned to the observations coming from 1 hidden state vs. 2 hidden states (Eq. 7) is shown as a function of amount of experience. Early in training, the two hypotheses have similar probabilities, whereas later 1 hidden state is overwhelmingly more probable. This corresponds to an increase in certainty over training, which would translate into a decreased tendency to remap, similar to the empirical observations. Note that the axis here has been flipped relative to Fig. 3C in order to match the axis of the empirical results shown in panel A.

We can model the dynamics of stabilization using a variant of the simulation described in the last section (Fig. 3B-C). The generative model in that simulation matches the experiment of Law et al. (2016) in that observations were drawn in an alternating fashion from two distinct distributions. At each observation, we take the log ratio of the probability that the current observation was drawn from the inferred alternating distributions that the previous observations were assumed to be drawn from (inferred blue distributions in Fig. 3B) compared to the probability of assigning to a novel state. The relative probability of assigning to a novel state gradually decreases over time, corresponding to the emergence of a “stable” map. Another way to understand map stabilization is to consider observations which are generated from a single distribution with mean 0. We can consider the same hypotheses as were considered in Fig. 3, namely, that there are either 1 or 2 hidden states. We consider the same hypotheses but the actual generative process has the opposite structure as Fig. 3. Early in learning, the partition evidence ratio is approximately indifferent between the one-state and two-state hypotheses, but gradually accumulates evidence in favor of the one-state hypothesis, corresponding to the emergence of a “stable” map. Indeed, early in learning, the animal does not know whether it is receiving observations from the simulation of Fig. 3B-C or the simulation of Fig. 4C-D, as they are indistinguishable. Only after extensive experience is the animal able to identify which generative process is generating its observations.

### Remapping due to non-Sensory changes

Remapping is not solely driven by sensory aspects of experience. For example, place fields can remap depending on internal variables such as movement direction or task (Smith and Mizumori, 2006a; Sanders et al., 2019; Wood et al., 2000; Muller et al., 1994). In general, it is known that place fields can remap depending on which direction the animal is running on a linear track (Markus et al., 1995; Battaglia et al., 2004). However, place fields tend not to remap based on running direction in an open field. This is most clearly shown in Markus et al. (1995). They compared two conditions, both of which occurred in an open field: one in which the animal was randomly foraging, and one in which the animal was running between four specific locations in one of two directions. They found that the extent of remapping in response to movement direction was larger in the directed foraging condition than in the random foraging condition (Fig. 5A) despite having the same sensory cues in the two conditions.

From the perspective of hidden state inference, we can draw an analogy with the remapping observed after training in the circle and square boxes (Fig. 3), replacing the sensory features of the environment with the non-sensory information about selfmotion. In the directed foraging case, observations are clearly separated into two states (clockwise movement and counterclockwise movement), whereas in the random foraging case, there is no consistent partition that could support the inference of multiple states.

We model this experiment in the following way. Again, we take observations to be 1-dimensional for simplicity, where the single feature is the animal’s movement direction. This feature is represented as a circular (angular) variable, as movement direction is circular. We model the random foraging condition as observations drawn from a uniform distribution over the circle (red dots in Fig. 5B). We model the directed foraging as observations drawn from a Von Mises distribution with *μ* = 0, *κ* = 10 alternating with a Von Mises distribution with *μ* = *π,κ* = 10 (blue dots in Fig. 5B). For each condition, we separate the observations into two groups with a line for which the distance from any observations is maximum (red and blue lines in Fig. 5B). After 10 observations, we ask the model what the relative probability is that the observations were drawn from a single hidden state or drawn from two hidden states split by the line of maximum separation. The model assigns greater probability to the two-state hypothesis for directed foraging. In contrast, it assigns greater probability to the one-state hypothesis for random foraging (Fig. 5C). This corresponds to the empirical finding that place fields were more likely to remap under the directed foraging condition compared to the random foraging condition.

### Morph experiments

A persistent puzzle in the field is the inconsistent results from “morph” environments that interpolate between different geometries (e.g., square and circle). Different labs have found different results with experimental setups that are not directly comparable (Wills et al., 2005; Leutgeb et al., 2005a; Colgin et al., 2010). We summarize the past results here and suggest an interpretation that leads to a novel prediction.

In 2005, two groups each performed an experiment to answer the question, “How does the hippocampus represent a novel environment that is intermediate between two familiar environments?” Both groups familiarized rats in square and circle environments, and then tested them in intermediate environments (polygons with a variable number of sides). The two papers had different results (Fig. 6A), characterized at the time in terms of whether the similarity curve had a discrete switch or a gradual switch. However, this difference is extremely hard to robustly characterize, considering that the variation in similarity between repetitions of the same environment was half as large as the entire range of similarity variations for the entire morph sequence (compare first and last points in Leutgeb et al., 2005a, their Fig. 6E)). The other difference that was discussed at the time was whether the population response was coherent or heterogeneous. While both studies showed heterogeneous population responses, they did show different levels of heterogeneity, and we discuss this below.

**Figure 5:**
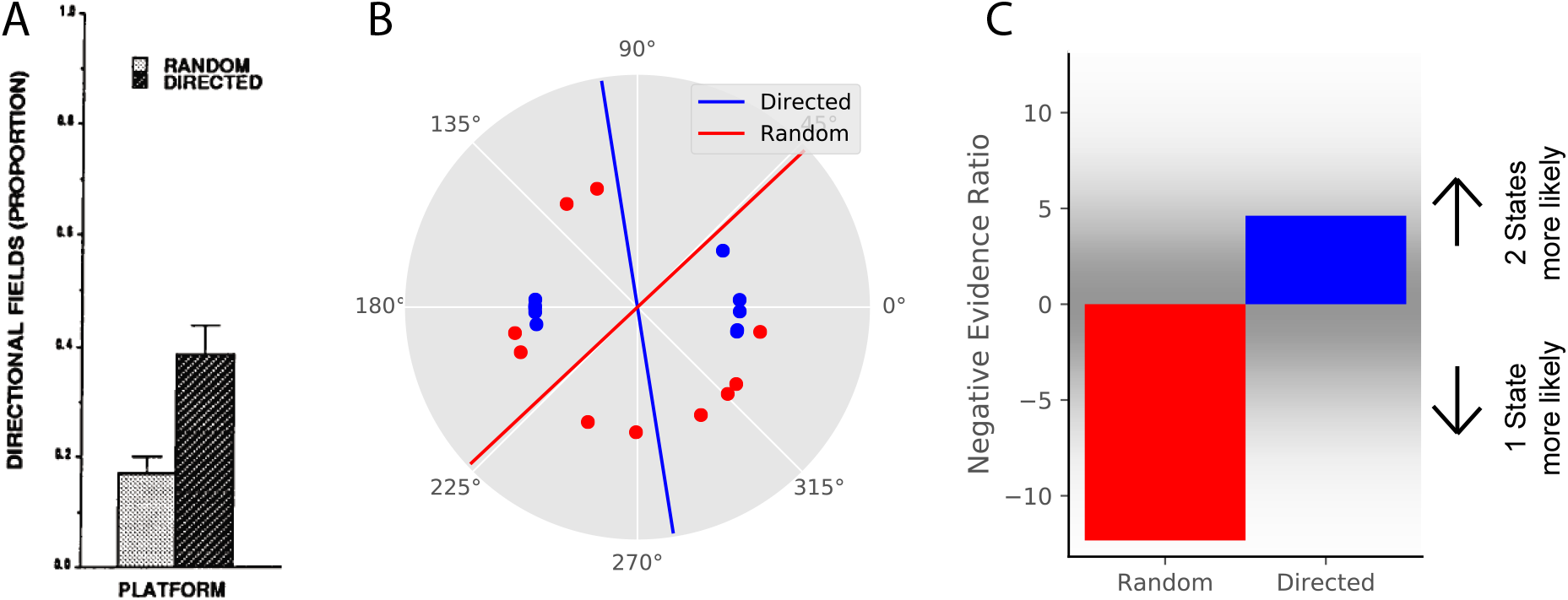
(A) Data from Markus, et al., 1995, showing that place field remapping depends on the animal’s direction more when the animal is running in a stereotyped path than when the animal is running in random directions. (B) The model receives circular observations corresponding to the animal’s running direction. The model either receives observations drawn from a uniform distribution (red dots) or alternating from two Von Mises distributions with means of 0 and 180 degrees, and *κ* = 10 (blue dots). These observations are separated into two groups with a line that is the farthest from any observations (red and blue lines). (C) The partition evidence ratio between the hypothesis that all observations having been drawn from two hidden states separated by the lines in panel B vs. having been drawn from a single hidden state (Eq. 7) after 10 observations. The model is more likely to put probability on the hypothesis that there are two hidden states when given the directional observations as opposed to the uniform observations. This is similar to the empirical results, where place fields are more likely to remap (more likely to infer two hidden states) when the animal is running in a directed fashion.

A much more striking point of comparison was the difference in the extent of remapping between the extreme square and circle environments. Complete remapping was observed between the square and circle in Wills et al. (2005), whereas partial remapping was observed in Leutgeb et al. (2005a). We believe that the findings of partial vs. complete remapping is the major difference in the findings of these papers, and is the one we focus on explaining.

What differences in protocols led to these differences in results? In addition to all the idiosyncrasies of individual lab protocols, there were two major explicitly described differences between their protocols. One is that they used different training protocols. Wills et al. (2005) used a training protocol designed for inducing complete remapping between square and circle in 6 days, and excluded animals that did not meet that criterion. Leutgeb et al. (2005a) used a similar training as Lever et al. (2002, see Fig. 3) for three weeks. The second difference was that Wills et al. (2005) presented the intermediate shapes in a scrambled order on the test day, whereas Leutgeb et al. (2005a) presented the intermediate shapes sequentially based on number of sides on the test day.

The second difference was the focus of several theoretical explanations (Blumenfeld et al., 2006; Gershman et al., 2014), but a replication of Wills et al. (2005) using scrambled presentation resulted in limited remapping (Colgin et al., 2010), demonstrating that a scrambled presentation was not sufficient to force the hippocampus to use complete coherent remapping. Differences in the training protocol remain as a possible explanation. However, the problem remains that Colgin et al. (2010) attempted an exact replication of Wills et al. (2005), but got the opposite result. These differences can be seen in Fig. 6A.

These results fit into a broader pattern of inconsistent results across two labs. Two experiments that led to complete remapping in the O’Keefe lab ended up leading to partial (and/or rate) remapping in the Moser lab. Training in alternating square and circle environments led to partial remapping initially and to complete remapping after 18 days in the O’Keefe lab (Lever et al., 2002), but led to partial remapping after 18 days of comparable training in the Moser lab (Leutgeb et al., 2005b,a). A 6 day white/morph circle-square training protocol led to complete remapping in the O’Keefe lab (Wills et al., 2005), but led to partial remapping in the Moser lab (Colgin et al., 2010). We do not believe either lab’s training to be inherently superior, but we do wish to point out that there are likely unreported idiosyncrasies of training that cause animals to consistently progress through partial remapping to global remapping more slowly in the Moser lab than in the O’Keefe lab (at least during the years 2000-2010). The main implication of this is that remapping behavior does not have a one-to-one mapping to the experimenter-defined conditions; rather, remapping behavior responds to a huge array of experiential factors, and the experimenter is only aware of a subset of these factors. Practically, this means that attempts to compare remapping behavior must be done between comparable controlled setups (as performed in the internal comparisons of Colgin et al., 2010), and comparisons should ideally not be made across labs.

**Figure 6:**
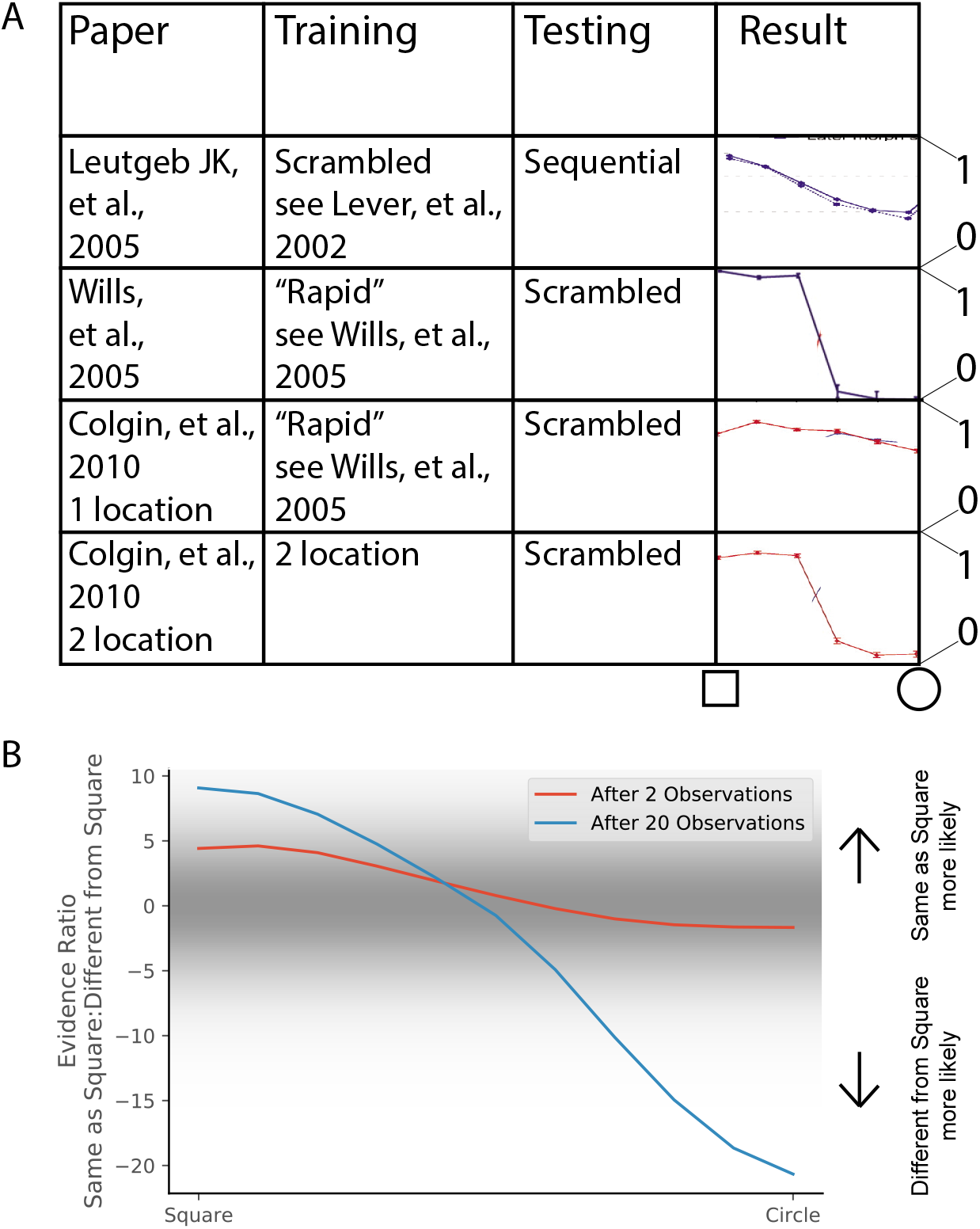
(A) Different experimental protocols give different results for the morph experiment. The results in the fourth column show the similarities in population representation of the intermediate morph shapes compared to the square shape. All values are shown on a scale ranging from 0 to 1, where 1 is complete concordance of population representations and 0 is random concordance. We classify the results into two qualitative classes: the first and third rows have results where all levels of morph result in partial remapping, whereas the second and fourth rows switch between no remapping and complete remapping as morph level increases. Scrambling during testing does not seem to be related to this effect. Moreover, the same experimental protocol can have qualitatively different results in different labs (compare second and third rows). (B) We provide observations from two alternating Gaussians with means −0.5 and +0.5, just as in Fig. 3. We test after 2 (red) or 20 (blue) training observations by providing intermediate values and measuring the relative probability of being assigned to the same hidden state as the −0.5 mean observations. We thus predict that both qualitative results can be achieved in the same lab simply by performing the morph testing at different points of training.

To summarize, various experimental protocols for measuring remapping behavior in response to intermediate “morph” environments give divergent results, which can be split into two categories: heterogeneous responses when there is only partial remapping between the extremes, and population-wide coherent responses when there is complete remapping between the extremes (Fig. 6A). As we explored above (Fig. 3), partial remapping and complete remapping can be observed in a single experimental protocol early and late in training, respectively. We therefore predict that both sets of results can be observed in the same lab, with the same experimental protocol, simply by presenting the intermediate “morph” environments early or late in training.

We show simulations of this prediction in Fig. 6B. Specifically, we compute the probability that the training observations came from a single hidden state *P*(**c1**) and the probability they came from two hidden states *P*(**c2**) according to Eq. 6. We then calculate the probability that the morph test is assigned to the same hidden state as the square assuming that the training observations came from two hidden states *P*(*c_probe_* = *c_square_*|**c2**) (Eq. 8). The hypotheses that correspond to the morph being assigned the same hidden state as the square are S1) that there is a single hidden state for the training and the morph is from the same state and S2) that there are two hidden states for the training and the morph is from the same state as the square. The hypotheses that correspond to the morph being assigned a different hidden state than the square are D1) that there is a single hidden state for the training and the morph is from a novel state and D2) that there are two hidden states for the training and the morph is from the same state as the circle and D3) that there are two hidden states for the training and the morph is from a novel state. We take the log posterior ratio between the S hypotheses and the D hypotheses and plot that in Fig. 6B for varying number of training experiences. The probability of assigning intermediate “morph” environments to the same hidden state as one of the extreme environments increases with the amount of training.

Thus, we suggest that a key distinction between classes of past morph results is whether there is complete or partial remapping between the extreme environments, and that complete or partial remapping can be achieved by a wide range of training protocols (as described throughout the paper) including amount of experience (as described in Fig. 3).

### Animal-to-animal variability

One challenge in the study of hippocampal remapping is that different animals respond differently to the same environments. Indeed, many of the previously discussed studies reported significant heterogeneity across animals in remapping behavior. Studies of the development of remapping over the course of learning frequently report that different animals learn at differing rates (Bostock et al., 1991; Lever et al., 2002). In fact, the variability across animals is frequently a nuisance in running experiments. The pre-training for one of the morph experiments described above (Wills et al., 2005) had three different ways that the observations could be partitioned. Out of the six animals they trained, four animals partitioned the observations in the way the experimenters expected, and the other two animals partitioned the observations in the other two possible ways (and therefore were excluded from the rest of the study).

The hidden state inference model offers one way to capture this heterogeneity across animals. The concentration parameter *α* (see Materials and Methods) controls the tendency to infer new hidden states when unexpected data are observed. Variation in this parameter was previously used to model age-dependent (Gershman et al., 2010, 2017) and individual (Gershman and Hartley, 2015) variability in learning. While partitioning large amounts of cleanly separated data is insensitive to changes of *α* over several orders of magnitude, *α* can have effects on partitioning of ambiguous or insufficient data. For example, if we take the learning of remapping explored in Fig. 3, changes in the value of α can alter the speed at which the model switches from preferring a one-state hypothesis to a two-state hypothesis (Fig. 7A). Moreover, if we take evidence ratios around 0 as indicative of partial remapping, different a values can lead to different lengths of time spent in the partial remapping regime, even for the exact same set of experiences.

To explore a second manifestation of animal variability, we ran a simulation resembling the training of Wills et al. (2005). We characterize observations with two features: shape and color of the enclosure. The white circle is characterized by a 2D Gaussian with means [0.5, 0.5], the morph circle is characterized by means [0.5, −0.5], and the morph square is characterized by means [−0.5, −0.5]; all standard deviations are 0.1. We provided the model with observations from these generative distributions according to the schedule used by Wills et al. (2005) (Fig. 7B). We then asked the model to assign an unnormalized posterior probability to the following hypotheses:

1. Each of the environments were drawn from separate hidden states (Fig. 7C, red bars), corresponding to “did not show wooden circle to morph-circle pattern transfer.”
2. The circles were the same and were different from the square (Fig. 7C, blue bars), corresponding to the selection criterion adopted by Wills et al. (2005).
3. All the experiences were drawn from a single hidden state (Fig. 7C, purple bars), corresponding to “failed to show rapid remapping in the morph-square and the wooden circle.”

Different values of *α* lead to variation in relative preferences for these hypotheses.

These results invite the interpretation that animal variability may be understood in terms of individual differences in the *α* parameter (though of course other parametric variations might produce some of the same effects).

### The effect of cue variability

In this section, we explore an experimental prediction of the model that highlights one of its key insights: remapping critically depends on past experience. Consider an environment that is characterized by two features. We can separate animals into two training groups: one in which feature 1 is highly variable and one in which feature 2 is highly variable (cyan and magenta dots in Fig 8A). We then probe with an observation that has a novel value in feature 1 (red x in Fig. 8A). The model predicts that an animal trained with higher variability in feature 1 will be more likely to assign the novel observation to the same state as the previous observations (i.e., not to remap; Fig. 8B). Intuitively, high variability will make the place fields more “tolerant” of deviations from the central tendency of the distribution.

By analogy, imagine a building with many similar conference rooms. One conference room always has its chairs arranged in a particular configuration (a low variability context), whereas another conference room frequently has different configurations (a high variability context). Intuitively, a change in the expected configuration in the low variability context will prompt the inference that you must be in a different room (and hence the place cells in your hippocampus will remap), whereas a change in the expected configuration in the high variability context will not. In the high variability context, you expect the unexpected (cf. the concept of “expected uncertainty” in Yu and Dayan, 2005).

**Figure 7:**
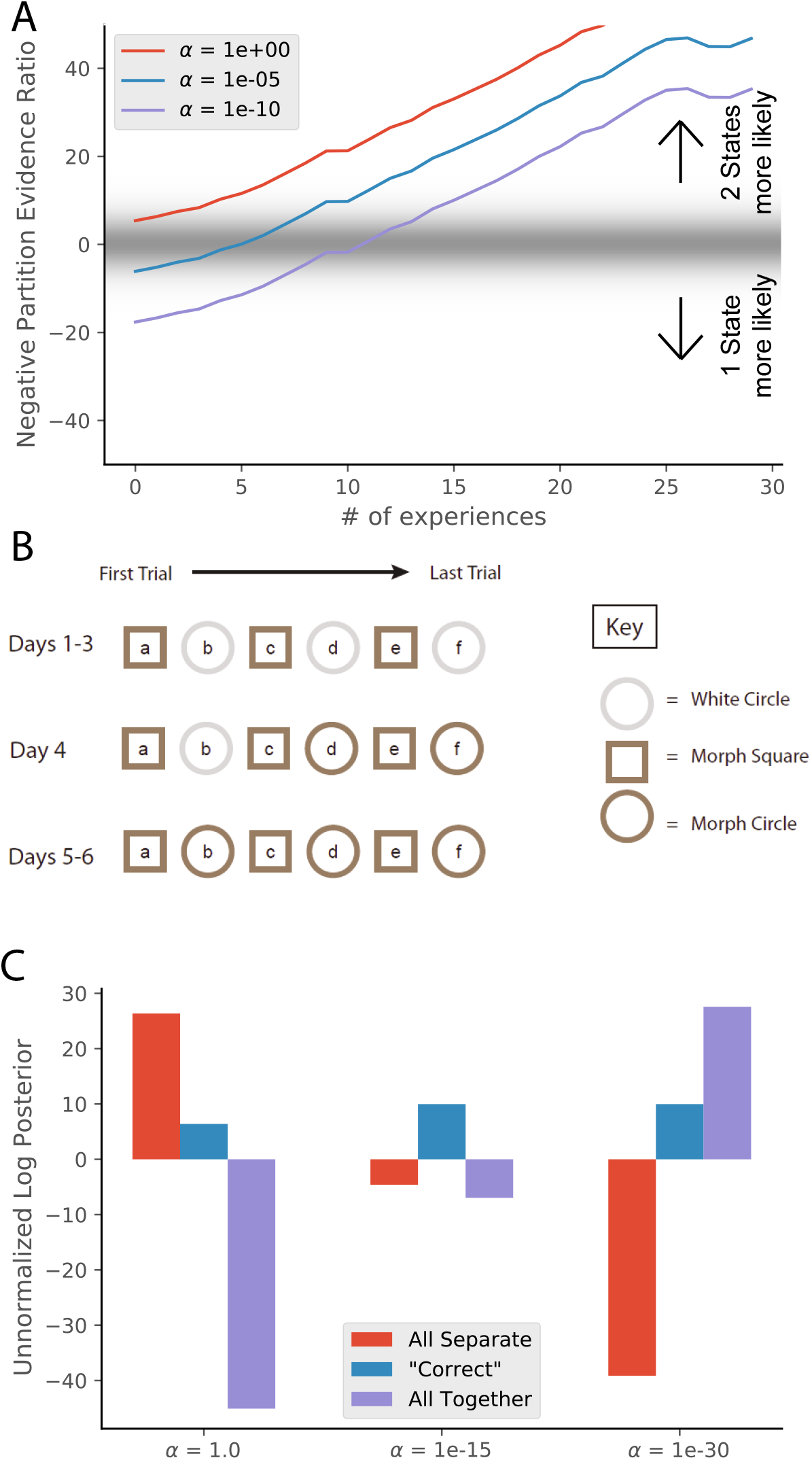
(A) Simulations from Fig. 3C with different values of α. Larger values of alpha lead to a greater tendency to infer a larger number of hidden states, and therefore a faster transition from preferring the single-state model to the two-state model. (B) The training protocol from Wills, et al. (2005). (C) In red is the probability assigned to the hypothesis that the white circle, morph circle, and morph square are all generated by separate hidden states. In blue is the probability assigned to the hypothesis that the white circle and morph circle are generated by the same hidden state and the morph square is generated by a separate hidden state, which is the hypothesis that the authors expected. In purple is the probability assigned to the hypothesis that all of the enclosures are generated by the same hidden state (Eq. 6). Different settings of α result in different preferred assignments of observations to hidden states, corresponding to the finding that different animals had different remapping behaviors.

**Figure 8:**
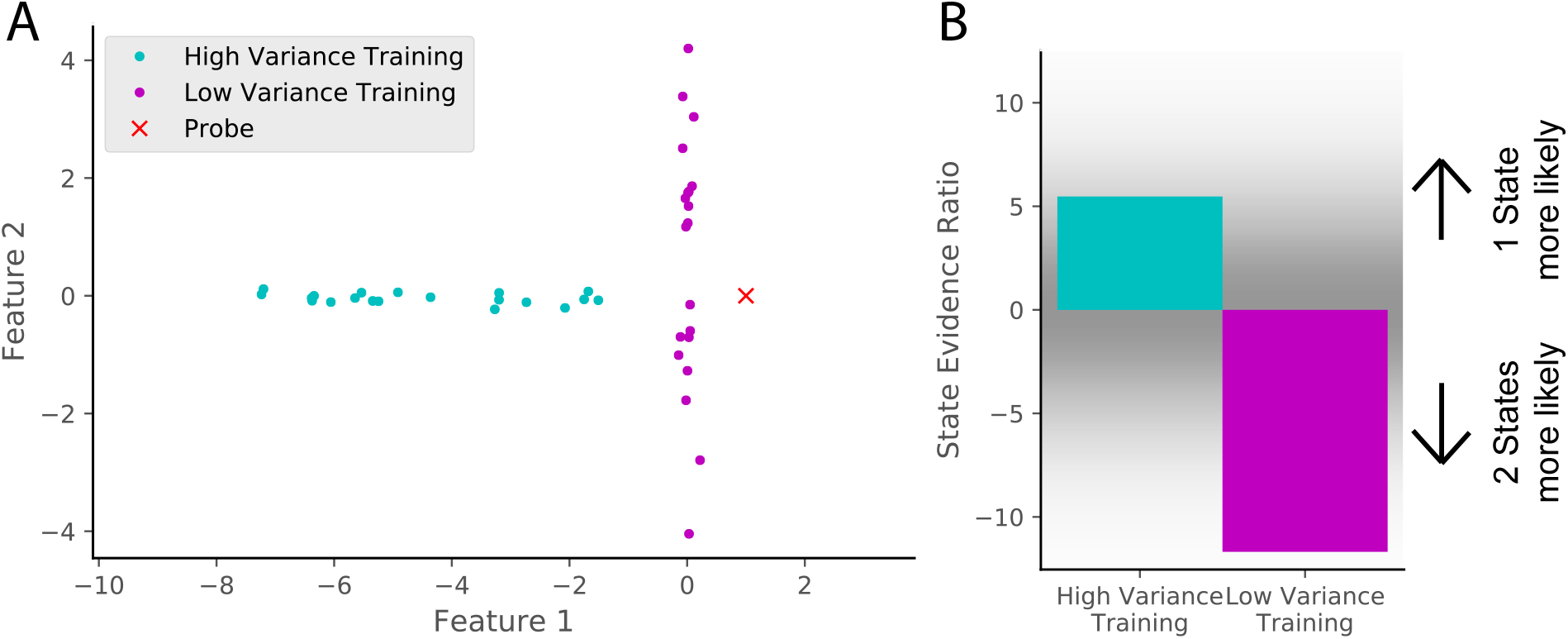
(A) Two training protocols (cyan and magenta) give (B) qualitatively different hidden state inferences when presented with the same novel observation (red dot in A). The cyan training is drawn from a Gaussian with mean [−5,0] and standard deviations [2, 0.1], whereas the magenta training is drawn from a Gaussian with mean [0,0] and standard deviations [0.1, 2]. The probe is presented after 20 training observations. The state evidence ratio here is the comparison between the assignment of the probe to the same hidden state as the training samples vs. a novel hidden state (Eq. 10).

## Discussion

We have proposed that hippocampal remapping provides a window into the process of hidden state inference. According to our framework, animals receive a stream of observations (data points), which they attempt to partition according to the hypothetical hidden states that generated them. Bayesian inference offers a natural solution to this problem. The specific form of Bayesian nonparametric model that we employed here has been previously invoked to explain a number of other hippocampal-dependent behavioral phenomena (Gershman et al., 2010, 2017, 2014). In this paper, we showed that this model recapitulates a broad range of remapping phenomena.

Central to our account is the idea that remapping reflects inferences about the hidden state, and in particular that partial remapping corresponds to high levels of uncertainty. Manipulations of sensory cues, environmental geometry, and training can all be understood in terms of their effects on state uncertainty. While this account has the potential to unify many phenomena under a common theoretical umbrella, there are still many loose ends and open questions, which we discuss below.

### What is the feature space?

Our model takes feature vectors as its inputs, but what are these features? In our simulations, we allowed them to be highly abstract idealizations. Ultimately, a biologically grounded theory must specify these features in terms of the inputs to the hippocampus. Furthermore, it will be necessary to more explicitly specify what timescale the model is operating on, since different features are relevant at different timescales. Although we have focused on the timescale of hours to days, map switches can occur on the subsecond timescale (Olypher et al., 2002; Jezek et al., 2011; Kelemen and Fenton, 2016).

One general hypothesis about the feature space encoded by the hippocampus is the *successor representation* theory (Stachenfeld et al., 2017), which posits that place cells encode a predictive map of the state space. On this view, the feature inputs to the hippocampus correspond to state features. This raises the intriguing possibility that remapping should be sensitive to predictive relationships between states. Many studies have observed that place cells are modulated by prospective information like the animal’s future trajectory (e.g., Battaglia et al., 2004; Ferbinteanu and Shapiro, 2003). It is less clear whether there is any evidence for global remapping as a function of changes in prospective information.

### Approximate inference

As discussed in the first section of the Results, exact inference over assignments of observations to hidden states is intractable, because the number of possible partitions is too large. As a result of this intractability, for most of the paper, we have limited ourselves to comparisons between a small number of hypotheses (selected based on the fact that most of the posterior probability will be concentrated on these hypotheses). This should be understood as an analytical heuristic rather than as an algorithmic theory of how the brain approximates probabilistic inference. A complete algorithmic theory must explain how the brain deals with arbitrarily large hypothesis spaces.

One idea is to model the hippocampus as stochastically sampling the hypothesis space (Fox and Prescott, 2010; Savin et al., 2014). According to this view, a sampling approximation approach would discretely represent each hypothesis with a frequency proportional to its probability. This fits nicely with the empirical finding that multiple maps can alternate rapidly (Kelemen and Fenton, 2016, 2010; Jackson and Redish, 2007; Kay et al., 2019; Jezek et al., 2011). Some of these findings suggest an oscillatory implementation, whereby each theta cycle plays the role of a single sample from the distribution of possible hidden states, and the extent of map switching corresponds to the degree of uncertainty about hidden state assignment. Indeed, map switching increases at points of uncertainty (Jezek et al., 2011). We would additionally predict that measures of map switching such as overdispersion would decrease over the course of experience in protocols such as that of Lever et al. (2002) as one hypothesis dominates (i.e., the evidence ratio between alternative hypotheses gets farther from 0 in Fig. 3C).

### Rotation experiments

There are several classes of empirical results that are related to the results explained in this paper, but not directly explained by our model. For example, in rotation experiments, the experimenters manipulated cues associated with the environment itself (“proximal cues” or “maze cues”) and/or manipulated cues associated with the room that the recording environment was placed in (“distal cues” or “room cues”). They asked questions such as whether the place cells followed the rotation of the maze cues or the room cues (Shapiro et al., 1997), and whether the place cells followed the animal’s own motion or the motion of the cues (Knierim et al., 1998). The answers to these questions were generally inconclusive, as they were sensitive to slight differences in protocol across labs. However, a consistent finding was that the results changed over the course of experience. For example, when a cue was repeatedly moved relative to other cues in an unstructured way, the cue lost control over the rotational alignment of the place fields (Knierim et al., 1995). While we do not explicitly model spatial relationships in our simulations, Knierim’s finding is similar to the training variance effect described in Fig 8: when the model is trained with observations for which a cue has high variance, further variation in that cue is less likely to cause a new hidden state to be inferred.

Conversely, in Shapiro et al. (1997), the maze cues and room cues were each rotated 90 degrees in opposite directions. Initially, the place cell representation split, some following room cues and some following maze cues. However, after a few repetitions, the place cell representation remapped between the two conditions. This is reminiscent of the simulations in Fig. 3, where a particular cue manipulation (square-circle) initially does not cause remapping, but after sufficient repetition, the place cells remap between the conditions; more evidence has been gathered to support the hypothesis that two distinct hidden states exist.

### Types of remapping

An influential interpretation of the literature has been that there are two main types of remapping: “global remapping” and “rate remapping”. In particular, it has been argued that global remapping corresponds to changes in physical location whereas rate remapping corresponds to changes in condition that occur at those locations (Leutgeb et al., 2005b; Colgin et al., 2010, 2008; Alme et al., 2014; Lisman et al., 2017). As discussed in the Introduction, the lines between global remapping and rate remapping are not so sharp. Global remapping can occur between conditions at the same physical location (Wills et al., 2005), and rate remapping can occur between different physical locations (Spiers et al., 2015). Moreover, the same manipulation can cause global remapping or rate remapping at different points in training (Lever et al., 2002). Our work provides an explanation for why there are not clear delineations of which manipulations cause which types of remapping. The animal must *infer* hidden states from its observations. Alternative hypotheses must be considered as long as ambiguity exists about the appropriate assignment of observations to hidden states. This uncertainty about hidden state assignment can manifest as “partial” or “rate” remapping. The statistics of these hidden states can be learned over the course of experience, leading to increased certainty about hidden state assignments. This increased certainty can be observed as more definitive “global” remapping or conversely, lack of remapping. One of our key points is that these categories are better thought of as existing along a continuum defined by state uncertainty.

### Relationship to other theories

How does this proposal relate to other theoretical perspectives on hippocampal remapping? We can contrast our model with a basic similarity threshold model, according to which each state is associated with a fixed set of features, and new observations would be classified as the same or different based on whether they exceed some threshold of change detection. This model does not capture some of the key phenomena associated with remapping; in particular, it cannot account for any of the ways in which learning affects remapping.

One major model of remapping is the attractor network. Based on early work by Hopfield (1982), the idea is that activity patterns associated with particular observations are learned by the network so as to be able to recover those activity patterns when degraded versions are presented. One attractor network implementation that has been specifically used to model remapping results was proposed by Blumenfeld et al. (2006). They sought to explain the difference in results between Wills et al. (2005) and Leutgeb et al. (2005a) by focusing on the scrambled order of the morph sequence. Their model was a conventional Hopfield network augmented with a “weight” term to change the pattern strength based on the novelty of that pattern. This led to attractors that were lumped together when the morph experiences were presented in sequential order instead of in a scrambled order. However, later work (Colgin et al., 2010) demonstrated that the order of presentation of the morph experiences was not the decisive factor in the qualitative results of the morph experiments (as described in more detail in the “Morph Experiments” section of the Results).

The attractor network perspective can be connected to our hidden state inference model by examining the probabilistic version of the Hopfield network, known as the *Boltzmann machine* (Ackley et al., 1985). The basins of attraction can be understood heuristically as feature configurations for distinct hidden states. One can make this heuristic connection more precise by defining an explicitly state-dependent energy function combined with a distribution over states, which would correspond to a mixture of Boltzmann machines (Nair and Hinton, 2009; Salakhutdinov et al., 2012).

In computational neuroscience, attractor networks are usually used as mechanistic descriptions of neuronal dynamics, unlike our hidden state inference model that operates at a higher level of abstraction. Thus, comparison of the two approaches is not entirely straightforward. It is possible that an attractor network could be used as an implementation of parts of the hidden state inference model. For example, inference about new states vs. old states is conceptually similar to the distinction between “pattern separation” in the dentate gyrus and “pattern completion” in CA3 (Knierim and Neunuebel, 2015; Rolls and Kesner, 2006). The attractor network describes *how* pattern separation and completion work. The hidden state inference model describes *why* pattern separation and completion work the way they do.

Our hidden state inference model is similar in spirit to the probabilistic model of remapping developed by Fuhs and Touretzky (2007). In that model, each context is represented by a Hidden Markov Model. Remapping is then formalized as a model comparison problem. Like our model, their calculation weighs both simplicity of a hypothetical partition and its fit with the observed data. They use their model to explain gradual remapping Lever et al. (2002), failure to generalize Hayman et al. (2003), and some aspects of reversal learning and sequence learning.

## Conclusion

Place field remapping has long been one of the most puzzling aspects hippocampal physiology, yet still lacks a comprehensive theoretical account. In this paper, we have taken steps towards such an account, starting with a normative formulation of the problem that we believe remapping is solving, namely hidden state inference. The algorithmic and biological underpinnings of this theory remain incomplete, setting a clear agenda for future theoretical work.

## Materials and Methods

### Generative model

We model the animal’s sensory inputs (observations) as a vector **y** = [*y*_1_,…, *y_D_*] consisting of *D* features. The specific representation of these features varies across experimental paradigms. The animal assumes that observations are generated by discrete hidden states. At each time point, a state is stochastically selected according to prior *P*(*c*), and the observation features are sampled from the observation distribution associated with that state, *P*(**y**|*c*, *θ_c_*), where *θ_c_* represents the parameters of the observation distribution for state *c*. For notational simplicity we will omit the time index *t* whenever it is unnecessary for the exposition.

We place a prior *P*(*θ_c_*) over the parameters and then marginalize to obtain the likelihood that a set of *m* observations **Y**_*c*_ = [**y**_1_,…, **y**_*t*_,…,**y**_*m*_] came from a single state *c*

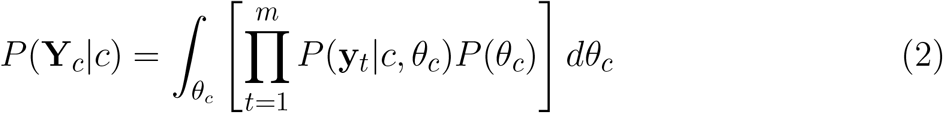

which we can extend to obtain the likelihood that a set of observations **Y** came from a set of *K* hidden states **c**

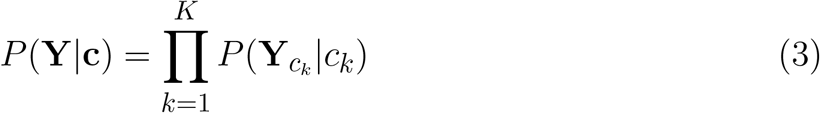

We model real-valued features with a multivariate normal observation distribution. The parameter vector is given by *θ_c_* = (*μ_c_*, Λ_*c*_), where *μ_c_* is the mean vector, and *Λ_c_* is the covariance matrix. We place a conjugate normal-Wishart distribution over these parameters (see Murphy, 2007, for more details), with hyperparameter values *μ*_0_ = 0 (prior mean), *κ*_0_ = 0.001 (scale parameter), *ν*_0_ = 0.02 (degrees of freedom), and *T*_0_ = 0.02 * *I* (scale matrix), where *I* is the *D*-dimensional identity matrix.

We model circular variables with a Von Mises observation distribution and a normal-gamma prior over the parameters. The hyperparamters of the prior are given by: *μ*_0_ = 0 (prior mean), *κ*_0_ = 0.001 (scale parameter), *α*_0_ = 0.01 (shape parameter), and *β*_0_ = 0.01 (rate parameter). Because in this case we cannot marginalize over parameters analytically, we used numerical integration.

To motivate our prior over hidden states, we start with a few basic desiderata: (i) the prior should be defined over an unbounded state space, allowing new states to be continually created; and (ii) the prior should prefer a small number of states, to facilitate generalization across observations (a form of Occam’s razor). These assumptions are satisfied by a simple nonparametric distribution known as the *Chinese restaurant process* (CRP; Aldous, 1985; Gershman and Blei, 2012), which samples states according to the following sequential process:

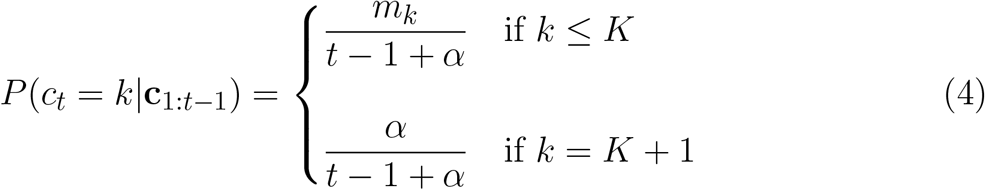

where *m_k_* is the number of previous observations assigned to state *k, K* is the total number of states created prior to time point *t*, and *α* ≥ 0 is a concentration parameter that controls the propensity to create new states. When *α* = 0, all observations will be generated by the same state. As *α* approaches infinity, each observations will be generated by a unique state. More generally, the expected number of states after *N* observations is *α* log *N*. Another way of using the CRP prior is to analytically calculate an unnormalized log probability to a given list of hidden state assignments **c**:

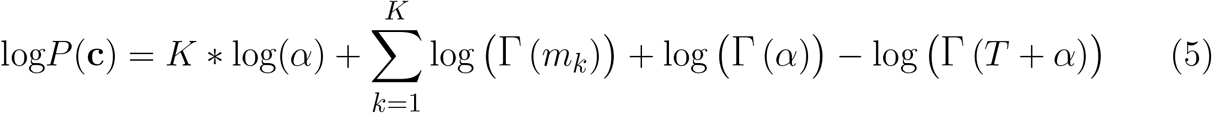

We set *α* = 0. 001 for all figures except Fig. 7, in which we explicitly explore the effects of variation in α. We emphasize that using this prior does not mean that the world actually generates hidden states through this process; it simply means that we are imputing this to the animal as its *internal model* of the world.

### Inference

To compute the posterior over hidden states, the likelihood is combined with a prior over state assignments, *P*(*c*), according to Bayes’ rule (Eq. 1). Because we are typically dealing with a set of observations, and hence a combinatorial space of state partitions (i.e., all possible assignments of observations to states), exact inference is intractable. However, because we are generally only interested in a small number of “plausible” partitions, we can simplify the problem by only assessing the relative probability of those states. The probability of each of those partitions c given a set of observations *Y*

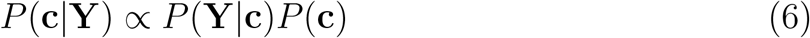

In particular, most of our simulations concern the question of whether two or three sets of observations are assigned to the same or different states. If we assume that all other partitions have probability close to 0, then we can ignore them without too much loss in accuracy. We use Eq. 6 in Fig. 7C.

The partition evidence ratio reported in the main text is the log odds ratio between the posterior probabilities of two hypotheses (partitions **c** and **c**’):

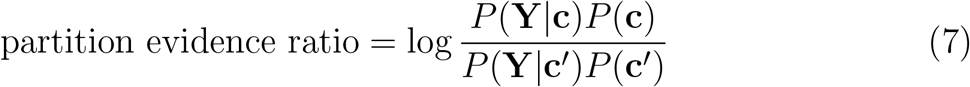

where **Y** denotes the set of observations, *P*(**Y**|**c**) is given by Eq. 3 and *P*(**c**) is given by Eq. 5. We use Eq. 7 in Figs. 3C, 4D, 5C, and 7A.

In some cases, we are interested in computing the posterior probability that a new observation *y*_*t*+1_ is assigned to a particular state conditional on a hypothetical assignment of all past observations:

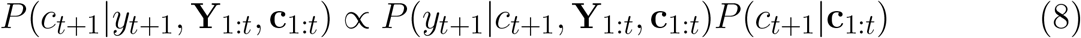

where *P*(*c*_*t*+1_|**c**_1:*t*_) is from Eq. 4 and *P*(*y*_*t*+1_|*c*_*t*+1_, **Y**_1:*t*_,**c**_1:*t*_) = *P*(*y*_*t*+1_|**Y***_c_k__*) is the posterior predictive distribution characterizing the probability of observing a value of *y*_*t*+1_ generated by a given hidden state *c_k_* given all previous observations **Y***_c_k__* with that hidden state assignment. For a Multivariate Normal likelihood function with a normal-Wishart prior, this is given by:

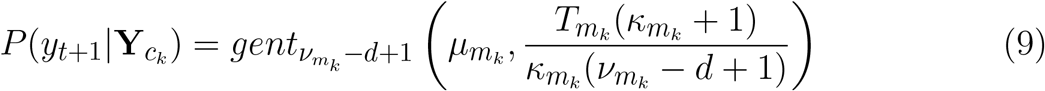

where *gent* is the generalized Student-t distribution with hyperparameters 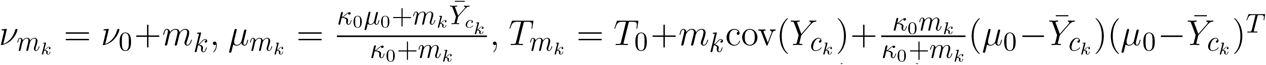, *κ_m_k__* = *κ*_0_ + *m_k_* as discussed in Section 8.3 of Murphy (2007).

The state evidence ratio reported in the main text is the log odds ratio between the posterior probabilities of two state assignments *c* and *c*’ for a given observation *y*_*t*+1_ given past state assignments **c**_1:*t*_ for past observations **Y**_1:*t*_

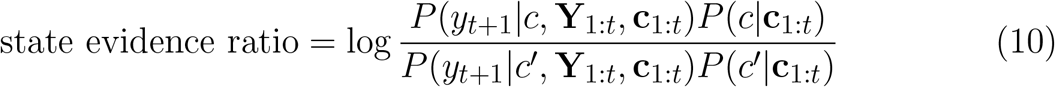

We use Eq. 10 in Figs. 1B-C,2, 4B, and 8B.

## Acknowledgments

We thank Hector Penagos for helpful comments on the manuscript. We thank Federico Claudi and Scidraw.io for the rat illustration in Fig. 1. This material is based upon work supported by the Center for Brains, Minds and Machines (CBMM), funded by NSF STC award CCF-1231216.

